# Iron oxidation is regulated by the two-component system, RegSR, and plays a role in photolithoheterotrophic growth in *Rhodopseudomonas palustris*

**DOI:** 10.1101/2021.08.19.456965

**Authors:** Nicholas W. Haas, Abhiney Jain, Zachary Hying, Sabrina J. Arif, Jeffrey A. Gralnick, Kathryn R. Fixen

**Affiliations:** BioTechnology Institute and Department of Plant and Microbial Biology, University of Minnesota, St. Paul, MN; Department of Genetics, Cell Biology, and Development, University of Minnesota, Minneapolis, MN

**Keywords:** iron oxidation, anoxygenic photosynthesis, *Rhodopseudomonas palustris*

## Abstract

Purple nonsulfur bacteria (PNSB) are metabolically versatile organisms generate energy through both aerobic and anaerobic respiration as well as anoxygenic photosynthesis. In many PNSB, the redox-sensing, two-component system RegBA is a global regulator of energy generating and consuming pathways, such as photosynthesis, carbon fixation, and nitrogen fixation, when cells are shifted from an aerobic to an anaerobic environment. However, in the PNSB *Rhodopseudomonas palustris,* the role of the RegBA homolog, RegSR, was unclear since global regulation of these same pathways involves the oxygen-sensing signal transduction system, FixJL-K, in *R. palustris*. Using RNA-seq analysis, we found that RegSR plays a role in regulating the operon *pioABC*, which encodes genes required for Fe(II) oxidation. We found that transcript levels of *pioABC* under photoheterotrophic conditions was dependent on the oxidation state of the carbon substrate and whether the cells were fixing nitrogen. We also found that *R. palustris* can carry out photolithoheterotrophic growth using Fe(II) oxidation when grown with the oxidized carbon substrate, malate, requiring *regSR* and *pioABC*. We present a model in which RegSR regulates *pioABC* in response to a cellular redox signal, allowing *R. palustris* to use Fe(II) oxidation to access more electrons when there is an increased cellular demand for reducing equivalents.

**Significance:** Mixotrophy is thought to be widespread in aquatic environments, yet little is understood about how mixotrophy affects biogeochemical cycles. Fe(II)-oxidizing anoxygenic phototrophs likely play an important role in iron cycling since they are thought to have thrived in the anoxic, iron-rich oceans of early Earth and can be found in both freshwater and marine environments. Although Fe(II) oxidation by anoxygenic phototrophs is largely studied in the context of photoautotrophic growth, these organisms can also grow photoheterotrophically. We present the first evidence linking photolithoheterotrophic growth using Fe(II) to the pathway required for photoautotrophic Fe(II) oxidation in an anoxygenic phototroph. Understanding this metabolism will be important for understanding how mixotrophic metabolism contributes to iron cycling in anoxic environments.

## Introduction

Bacterial metabolism is often studied in the context of a single nutritional mode, which does not reflect the spectrum that can exist between different trophic modes such as autotrophy versus heterotrophy. Mixotrophy often refers to an organism’s ability to use, sometimes simultaneously, different trophic modes for acquisition of carbon or energy, but it can also encompass acquisition of nitrogen, sulfur, and other trace elements. Although mixotrophy is thought to predominate in certain environments (1–4), little is understood about how mixotrophic organisms contribute to biogeochemical cycles. This is particularly true for iron cycling since microorganisms that harness the one-electron cycling between Fe(II) and Fe(III) play a major role in global biogeochemical cycling of iron, particularly in microaerobic and anoxic environments (5–10). Fe(II) is readily oxidized to Fe(III) in the presence of oxygen, which means in microaerobic and anoxic environments, such as the deep sea or aquatic sediments, Fe(II) oxidation is largely due to microbial activity. Lithoheterotrophic growth using Fe(II) oxidation has been engineered in an obligate aerobic chemolithoautotroph (11), and mixotrophy has been proposed for nitrate reducing and phototrophic Fe(II) oxidizers (12–18). However, for many of the Fe(II)-oxidizers that have been proposed to carry out mixotrophic growth, it is still unclear if Fe(II) oxidation is linked to enzymatic activity or is the result of abiotic factors.

Mixotrophic growth using Fe(II) oxidation may be particularly prevalent in Fe(II)-oxidizing anoxygenic phototrophs, which are widely distributed in both freshwater and marine environments (9). This group uses anoxygenic photosynthesis to obtain energy from light and can carry out photoautotrophic growth using Fe(II) as an electron donor for carbon dioxide (CO_2_) fixation in a type of metabolism referred to as photoferrotrophy (19, 20). All isolated Fe(II)-oxidizing anoxygenic phototrophs are metabolically versatile, capable of growing photoautotrophically or photoheterotrophically (16, 19–26). Although Fe(II) oxidation is largely thought to be involved in photoautotrophic growth, some anoxygenic phototrophs oxidize Fe(II) in the presence of an organic carbon substrate while others can oxidize Fe(II) but cannot use Fe(II) as the sole electron donor for growth (13, 16, 18). A role for Fe(II) oxidation as a detoxification mechanism has been proposed (25), but in most cases, the reason why some anoxygenic phototrophs oxidize Fe(II) under photoheterotrophic conditions remains unknown.

Most of our understanding of photoferrotrophy comes from work done in the purple nonsulfur bacterium (PNSB) *Rhodopseudomonas palustris*. *R. palustris* grows best as a photoheterotroph using energy generated by cyclic photophosphorylation and by metabolizing organic acids like malate or succinate and is known for its ability to fix nitrogen (27, 28). *R. palustris* can also grow as a photoautotroph using Fe(II), hydrogen, or thiosulfate as an electron donor (21, 29–31). Iron oxidation in *R. palustris* TIE-1 requires the *pioABC* operon (20), which encodes PioA, a decaheme *c*-type cytochrome; PioB, an outer membrane porin; and PioC, a periplasmic high-potential iron protein (HiPIP) (33, 34). These proteins are thought to form an electron transfer pathway that carries electrons from oxidation of Fe(II) at the outer membrane to the quinone pool in the inner membrane and are used to reduce NAD^+^ via reverse electron transfer (35). Although this operon is conserved in 34 of the 40 sequenced *R. palustris* strains, even closely related strains can vary in their ability to carry out photoferrotrophy. *R. palustris* CGA010, which shares an average nucleotide identity >97% with TIE-1, encodes *pioABC* but does not grow photoautotrophically using Fe(II) as the electron donor (32).

In TIE-1, expression of *pioABC* is induced under anaerobic conditions in both photoheterotrophic and photoautotrophic conditions (36). Induction of *pioABC* under anaerobic conditions requires the global regulator FixK (36). FixK is part of the oxygen-sensing signal transduction pathway FixJL-K and regulates expression of genes involved in photosynthesis, electron transfer, carbon fixation, aromatic compound degradation, and iron oxidation in response to a shift from aerobic to microaerobic or anaerobic conditions (36, 37). It is unclear why *pioABC* is expressed under non-photoferrotrophic conditions, but *R. palustris* TIE-1 has been shown to oxidize Fe(II) while growing photoheterotrophically on lactate but not acetate (13). This suggests that Fe(II) oxidation could play a role in photoheterotrophic growth, but it is unclear if *pioABC* is required for Fe(II) oxidation under photoheterotrophic conditions, and why Fe(II) oxidation occurs when lactate is consumed but not acetate (13).

In this study, we initially set out to understand the role of RegSR, a homolog of the redox-sensing, two-component signal transduction system, RegBA. We found that RegSR plays a role in regulating expression of *pioABC* under photoheterotrophic conditions, and expression was dependent on the oxidation state of the carbon substrate provided. Furthermore, we present the first evidence of a role for Fe(II) oxidation by PioABC in photolithoheterotrophic growth on an oxidized carbon substrate. These findings indicate that Fe(II) oxidation is not only important for photoautotrophy but also plays a role in photomixotrophic growth, in which Fe(II) acts as an additional electron donor to augment photoheterotrophic growth on an oxidized carbon substrate. A role for Fe(II) oxidation in photolithoheterotrophic growth may explain the presence of Fe(II) oxidation genes in PNSB that are unable grow photoautotrophically using Fe(II) as the electron donor.

## Results

### RegSR regulates expression of *pioABC* and plays a role in iron oxidation in *R. palustris* CGA010

In other PNSB, RegBA responds to changes in the redox state of the cytoplasm as well as the redox state of the quinone pool and acts as a global regulator of pathways that include photosynthesis, nitrogen fixation, hydrogen uptake, carbon fixation, and electron transport in *R. capsulatus* and *R. sphaeroides* (38–42). However, unlike other PNSB, the role of RegSR, a homolog of RegBA, in *R. palustris* was largely unclear. In *R. palustris,* global regulation of photosynthesis, carbon metabolism, and electron transfer (including *pioABC*) in response to changes in oxygen tension is controlled by FixJL-K, not RegSR, and *R. palustris* is one of the few PNSB shown to encode both FixJL-K and RegSR (36, 37). Although growth rates were not reported, it was observed that a *ΔregSR* mutant (CGA2023) grew similar to wild-type *R. palustris* (CGA010) under photoheterotrophic, photoautotrophic, and nitrogen-fixing conditions, which suggested that RegSR does not regulate photosynthesis, nitrogen fixation, or CO_2_ fixation in *R. palustris* (29, 37). In agreement with this observation, we found that CGA2023 showed only a slight defect in growth compared to CGA010 when grown photoheterotrophically, photoautotrophically, or under nitrogen-fixing conditions (Figure S1). This is consistent with FixJL- K acting as the global regulator of photosynthesis and nitrogen fixation in *R. palustris*, not RegSR.

To gain insight into the role of RegSR in *R. palustris*, RNA-seq was used to determine gene expression changes between CGA010 and CGA2023 under photoheterotrophic conditions. RegSR has been reported to play a role in expression of the gene, *hupS*, which encodes a subunit of an uptake hydrogenase in CGA010 (29). RegSR repressed expression of *hupS*, and repression of *hupS* was influenced by factors that affect the cellular redox status. These factors included conditions that require nitrogen fixation, a process known to consume reducing equivalents, and the oxidation state of the carbon source provided (29). To consider the effect that cellular redox status has on RegSR-dependent gene expression, CGA010 and CGA2023 were grown under non-nitrogen-fixing or nitrogen-fixing conditions and provided carbon substrates with different oxidation states– malate, succinate, acetate, or benzoate (Table 1).

**Table 1.** Oxidation states of *R. palustris* biomass and growth substrates

| Compound | Formula | Oxidation state<sup>a</sup> |
| --- | --- | --- |
| Malate | C<sub>4</sub>H<sub>6</sub>O<sub>5</sub> | +1 |
| Succinate | C<sub>4</sub>H<sub>6</sub>O<sub>4</sub> | +0.5 |
| Acetate | C<sub>2</sub>H<sub>4</sub>O<sub>2</sub> | 0 |
| Benzoate | C<sub>7</sub>H<sub>6</sub>O<sub>2</sub> | -0.3 |
| Biomass<sup>b</sup> | CH<sub>1.8</sub>N<sub>0.18</sub>O<sub>0.38</sub> | -0.5 |

<sup>a</sup> calculated as described in (56).579 <sup>b</sup> as reported in (56) and based on elemental composition of *R. palustris* 420L.

When we compared gene expression changes between CGA010 and CGA2023, we were surprised to find that one of the biggest changes in gene expression was for the operon involved in iron oxidation, *pioABC*. Expression levels of this operon were almost 20-fold less in CGA2023 grown photoheterotrophically with malate (Fig. 1). This change in gene expression was also dependent on the oxidation state of the carbon source provided and whether the cells were fixing nitrogen (Fig. 1). Expression of this operon decreased when a carbon source more reduced than malate was provided and expression was highest under nitrogen-fixing conditions (Fig. 1 and Fig. S2). This indicates that expression of *pioABC* is correlated with cellular demand for reducing equivalents, and RegSR is required for increased *pioABC* expression under these conditions.

**Figure 1.**
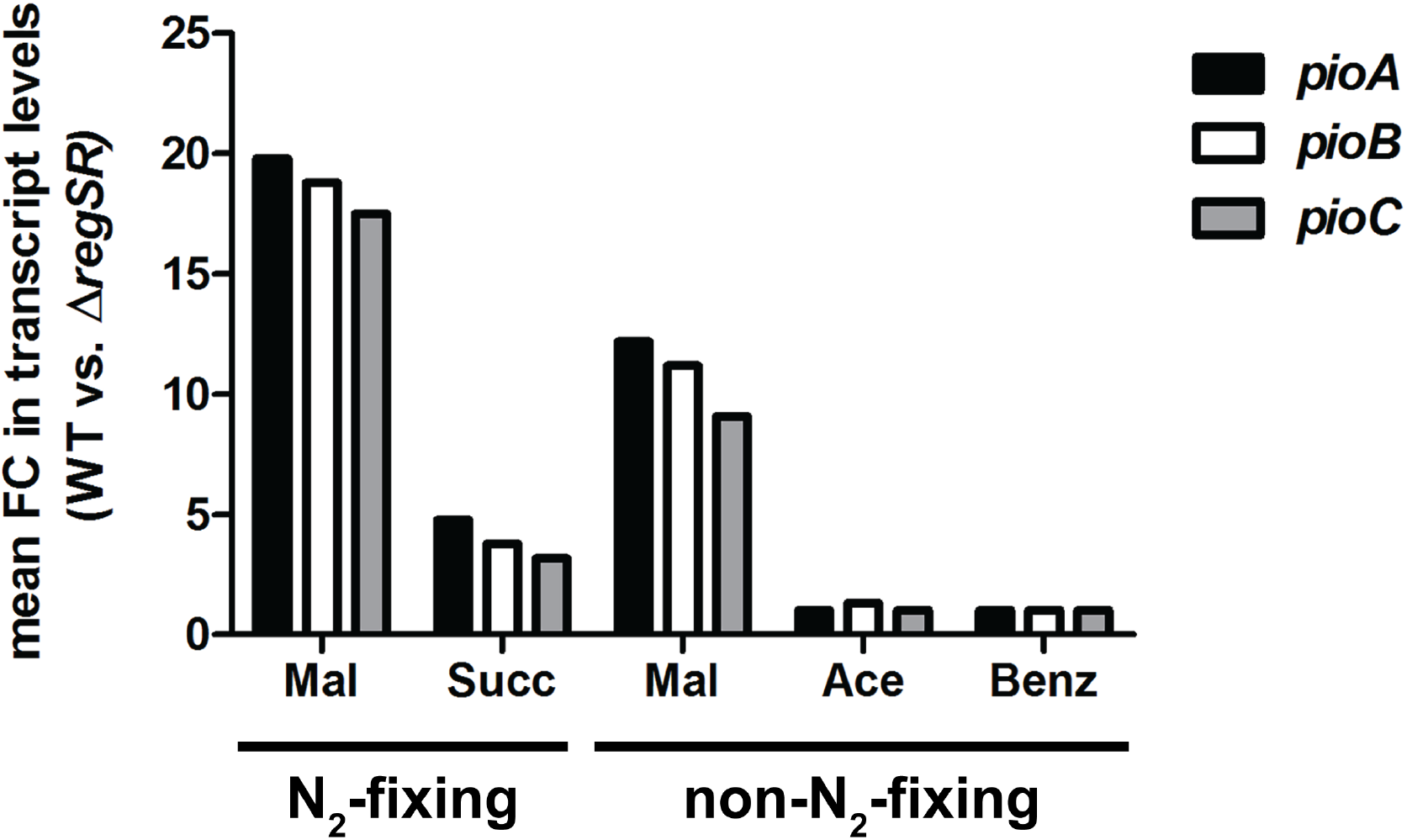
Expression of *pioABC* requires RegSR and is affected by both the oxidation state of the carbon substrate and nitrogen fixation under photoheterotrophic conditions. Shown is the fold change (FC) in *pioABC* transcript levels in *R. palustris* CGA010 (WT) versus CGA2023 (Δ*regSR*) as determined by RNA-seq analysis. Data are averages from duplicate cultures. Cells were grown photoheterotrophically using the indicated carbon substrate, supplied at a final concentration of 40 mM carbon, with (non-N_2_-fixing) or without (N_2_-fixing) ammonium sulphate. Carbon substrates are ordered by decreasing oxidation state. Mal, malate; Succ, succinate; Ace, acetate; Benz, benzoate.

Since RegSR regulates expression of *pioABC*, the ability of CGA2023 to carry out iron oxidation was tested. *R. palustris* CGA010, the parent strain of CGA2023, is not able to grow photoautotrophically using Fe(II). However, CGA010 grown photoautotrophically with thiosulfate or H_2_ can oxidize Fe(II), and its ability to carry out iron oxidation is dependent on an intact *pioABC* operon (32). To determine if RegSR plays a role in iron oxidation, we used a ferrozine assay to measure Fe(II) oxidation by cell suspensions of CGA010 and CGA2023 grown photoautotrophically with thiosulfate as the electron source. As shown in Fig. 2, CGA2023 was defective in its ability to carry out iron oxidation when compared to CGA010. Furthermore, complementation of *regSR in trans* in CGA2023 showed increased Fe(II) oxidation compared to CGA2023 with the vector alone (Fig. 2). This indicates that RegSR regulates Fe(II) oxidation, and this is likely because RegSR regulates expression of *pioABC*.

**Figure 2.**
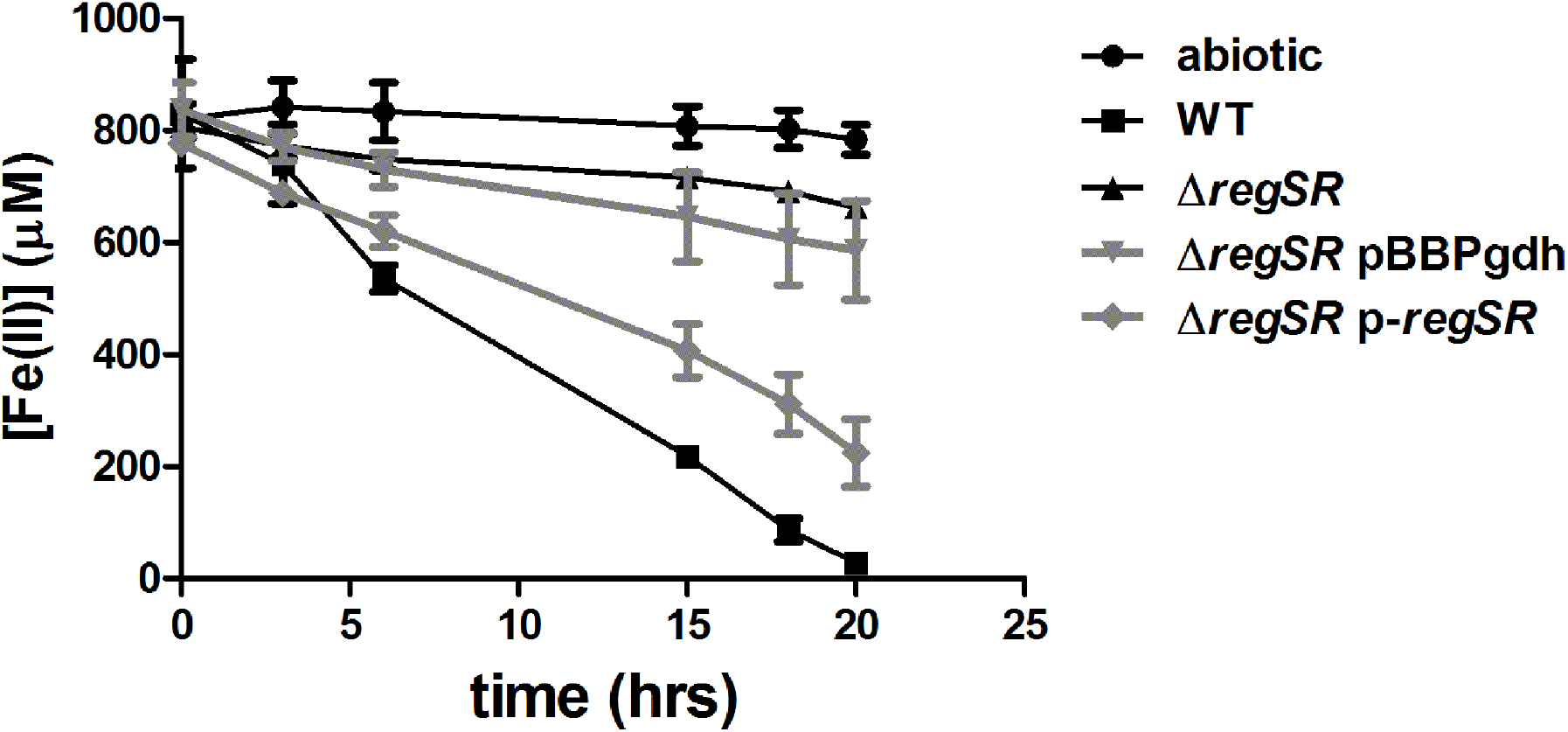
RegSR is required for Fe(II) oxidation by *R. palustris* CGA010. Fe(II) quantification over time for *R. palustris* CGA010 (WT), CGA2023 (*ΔregSR*), CGA2023 containing the expression vector pBBPgdh, and CGA2023 with *regSR* expressed *in trans* (p-*regSR*) cell suspensions. Cells were grown photoautotrophically with thiosulfate and CO_2_. Fe(II) was also quantified over time for samples without *R. palustris* cells (abiotic). Data represent the average of three replicates and the error bars represent standard deviation.

### *R. palustris* is capable of photolithoheterotrophic growth using Fe(II)

From the data presented above, RegSR plays a role in regulating *pioABC* expression and plays a role in iron oxidation. However, it was unclear why *pioABC* is expressed under photoheterotrophic conditions, and why its expression increases when an oxidized carbon substrate is provided. *R. palustris* has been shown to oxidize thiosulfate or H_2_ during photoheterotrophic growth on an oxidized carbon substrate, particularly when these substrates are limiting (less than 40 mM total carbon) (29, 31, 43). This allows *R. palustris* access to an additional electron source to allow complete assimilation of oxidized carbon compounds (29, 31, 43). We hypothesized that like thiosulfate and H_2_ oxidation, Fe(II) oxidation may provide an additional source of electrons, leading to increased cell yields under photoheterotrophic conditions with an oxidized carbon substrate.

To test if cells grown photoheterotrophically can carry out Fe(II) oxidation, we measured Fe(II) oxidation using a ferrozine assay with CGA010 grown with acetate or malate as a carbon substrate in the presence of 2.5 mM of Fe(II). Both organic acids were limited to a final concentration of 20 mM total carbon (e.g., 5 mM malate and 10 mM acetate). As shown in Fig. 3, cells growing photoheterotrophically with acetate did not oxidize Fe(II), and Fe(II) oxidation was comparable to an abiotic control. However, cells grown photoheterotrophically with malate oxidized Fe(II), indicating that Fe(II) oxidation can occur under photoheterotrophic conditions but only when malate is provided (Fig. 3).

**Figure 3.**
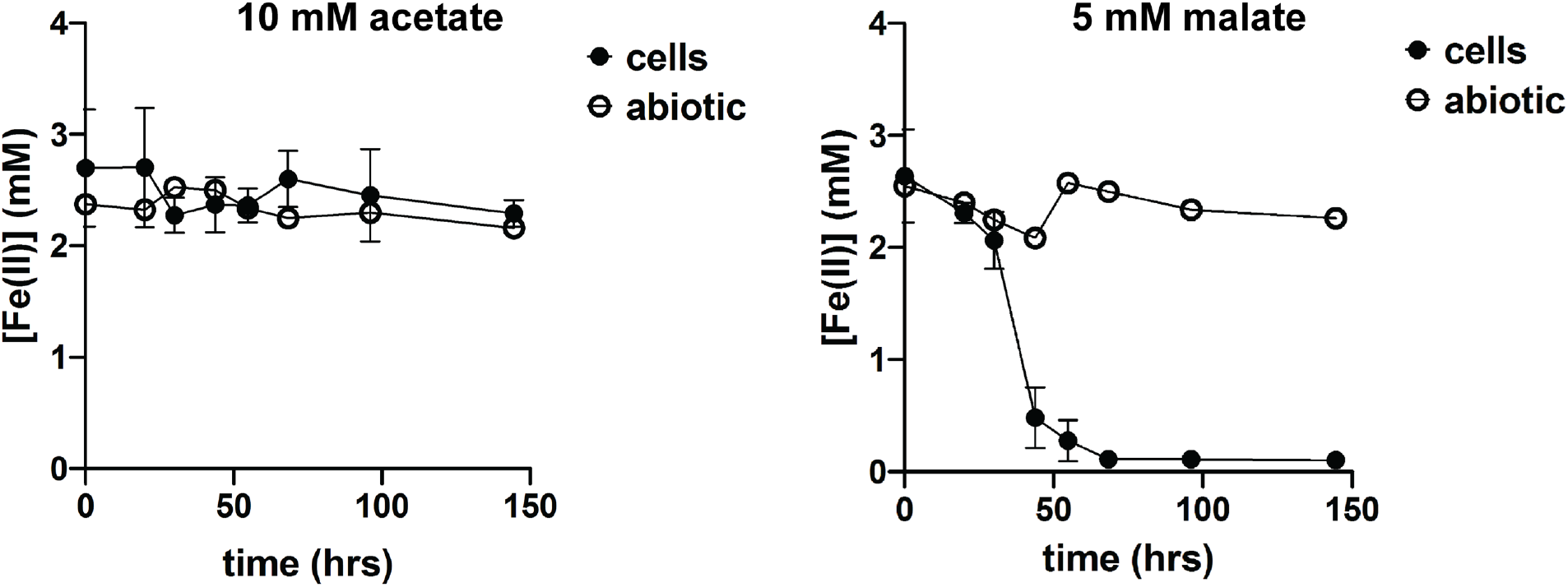
Photolithoheterotrophic growth using Fe(II) by *R. palustris* is dependent on the carbon substrate provided. Fe(II) quantification over time for *R. palustris* CGA010 (WT) growing in minimal medium lacking ammonium sulfate with 2.5 mM FeCl_2_, 2.5 mM nitriloacetic acid, and either 10 mM acetate or 5 mM malate (20 mM carbon total). Fe(II) was also quantified over time for samples without *R. palustris* cells (abiotic). Data represent the average of three replicates and the error bars represent standard deviation.

Since thiosulfate and H_2_ oxidation was shown to increase cell yield on oxidized carbon substrates, we looked at whether or not Fe(II) oxidation affected cell yield of CGA010 grown with malate or acetate as a carbon source. When provided with 2.5 mM Fe(II), cells grew to a higher optical density and had a higher total protein yield than cells grown in medium with a low concentration of Fe(II) (2.5 μM) (Fig. 4 and Fig. S2). However, this difference, although significant, was small when cells were grown under non-nitrogen-fixing conditions, and this difference increased under nitrogen-fixing conditions, resulting in an almost 1.5-fold increase in total protein yield (Fig. 4A). This difference in cell density and protein yield was not observed when cells were grown with acetate (Fig. 4B). The increase in cell density and protein yield also required RegSR since CGA2023 did not show a significant increase in cell density or protein yield when in the presence of a high concentration of Fe(II), even when grown under nitrogen-fixing conditions (Fig. 4 and Fig. S3). This indicates that Fe(II) oxidation can play a role in photolithoheterotrophic growth in which light serves as the energy source, inorganic compounds are used as electron donors, and organic compounds are consumed for cellular carbon (44).

**Figure 4.**
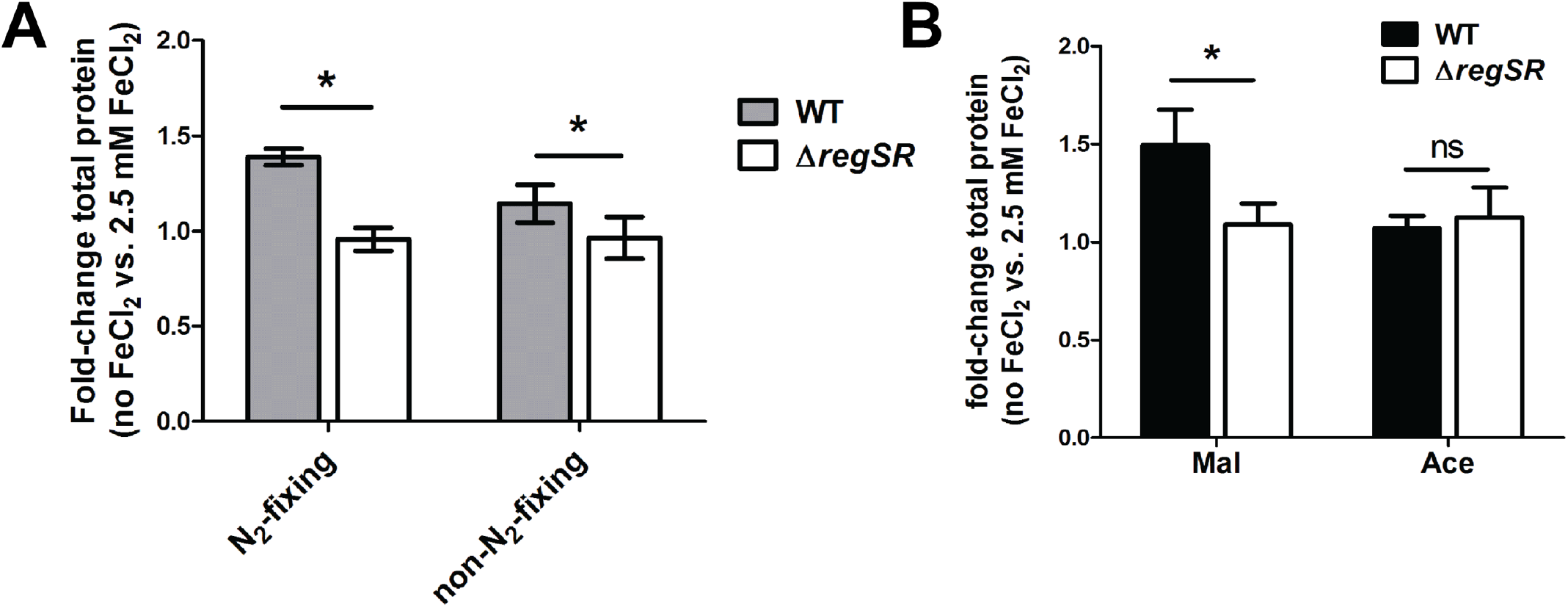
*R. palustris* cell yield increases under Fe(II)-oxidizing conditions. (A) Total protein was determined for stationary phase *R. palustris* CGA010 (WT) and CGA2023 (Δ*regSR*) cultures grown in minimal medium with (non-N_2_-fixing) or without (N_2_-fixing) ammonium sulfate and 5 mM malate. (B) Total protein was determined for stationary phase *R. palustris* CGA010 (WT) and CGA2023 (Δ*regSR*) cultures grown in minimal medium without (N_2_-fixing) ammonium sulfate and 5 mM malate or 10 mM acetate (20 mM total carbon). Both (A) and (B) show the ratio of total protein determined for cultures with 2.5 mM FeCl_2_ versus the total protein for culture without added FeCl_2_. Data represent the average of three replicates and the error bars represent standard deviation. Significance was determined using a two-tailed paired t-test, and \**p*<0.05. ns, not significant

To determine if *pioABC* is required for photolithoheterotrophic growth in *R. palustris*, in-frame deletion mutants in *pioA* and the *pio* operon (*pioABC*) were generated and tested for their ability to carry out Fe(II) oxidation during photoheterotrophic growth with malate. As shown in Fig. 5A, like CGA2023, cells containing a deletion in *pioA* or *pioABC* were unable to carry out Fe(II) oxidation. An intact *pio* operon was also required to see an increase in cell density and protein yield when grown with 2.5 mM Fe(II) (Fig. 5B and Fig. S3). While CGA010 resulted in an almost 1.5-fold increase in protein yield, CGA2023 and *R. palustris* cells containing a deletion in *pioA* or the entire *pio* operon did not show a significant change in protein yield (Fig. 5B). This indicates that PioABC is required for photolithoheterotrophic growth using Fe(II) oxidation.

**Figure 5.**
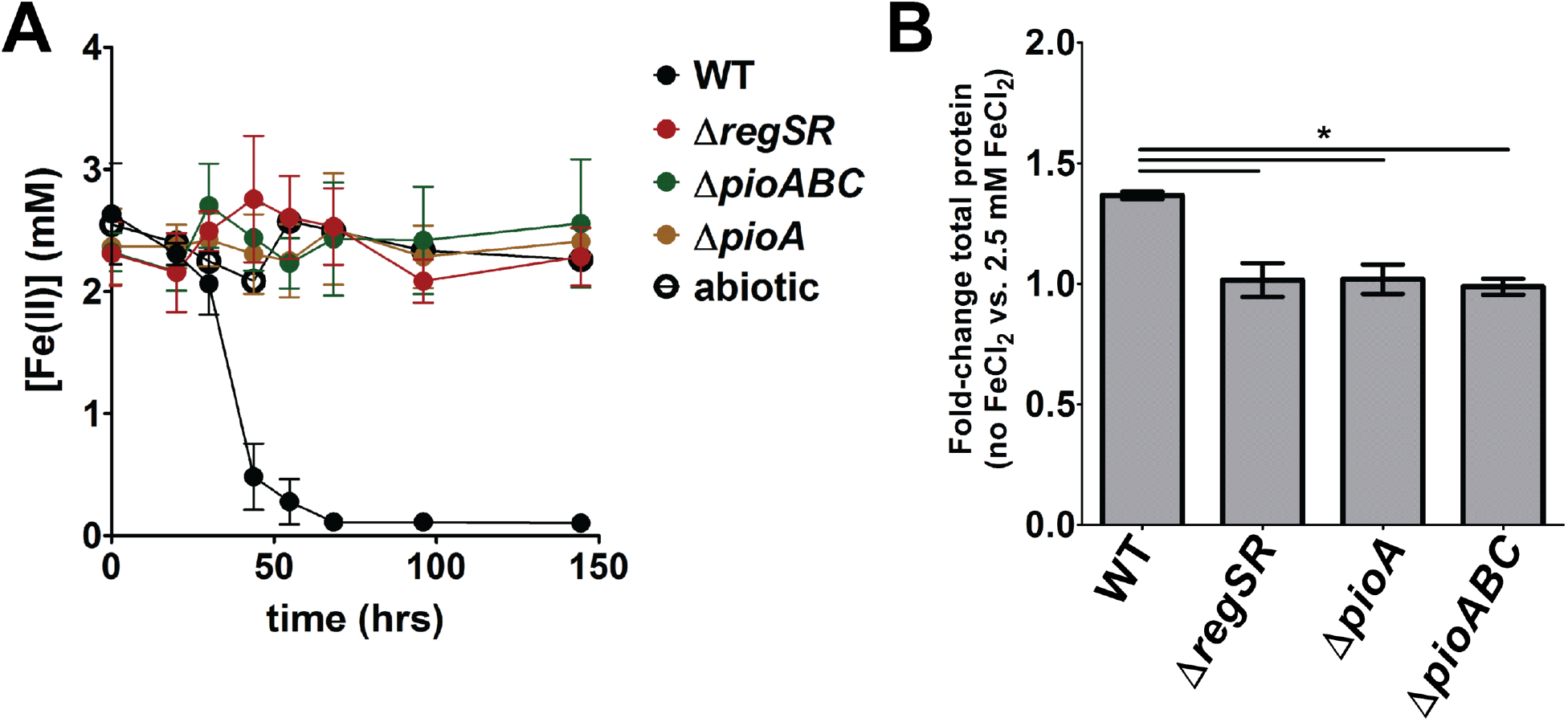
Photolithoheterotrophic growth using Fe(II) by *R. palustris* requires *pio* genes. (A) Fe(II) quantification over time for *R. palustris* CGA010 (WT) and CGA2023 (Δ*regSR*), Δ*pioA*, and Δ*pioABC* strains growing in minimal medium lacking ammonium sulfate with 2.5 mM FeCl_2_, 2.5 mM nitriloacetic acid, and 5 mM malate. Fe(II) was also quantified over time for samples without *R. palustris* cells (abiotic). (B) Total protein was determined for stationary phase *R. palustris* CGA010 (WT) and CGA2023 (Δ*regSR*), Δ*pioA*, and Δ*pioABC* cultures grown in minimal medium without ammonium sulfate, 5 mM malate, and with or without 2.5 mM FeCl_2_ added. Shown is the ratio of total protein determined for cultures with 2.5 mM FeCl_2_ added versus the total protein for culture without added FeCl_2_. Data represent the average of three replicates and the error bars represent standard deviation. \**p*<0.05 using a two-tailed, paired t-test.

### Other genes regulated by RegSR

As shown in Table S1, in all the conditions tested, 16 genes were more highly expressed in CGA2023 than CGA010, and 52 genes were down-regulated in CGA2023 versus CGA010, which is consistent with RegR and its homologues acting predominantly as transcription activators. Of the 68 genes that were differentially expressed, 35 percent (24 genes) encoded proteins of unknown function. Of the remaining genes, two genes, *phaA* and *phaC,* encode components of a putative potassium efflux system for adaptation to high pH. Both of these genes were down-regulated under all conditions (Table S1). To test if CGA2023 was more sensitive to pH, CGA010 and CGA2023 were grown at pH 6, pH 7, and pH 8. As shown in Fig. S4, CGA2023 grew like CGA010 in pH 6 and pH 7 but was more sensitive to high pH, which correlates with decreased expression of *phaA* and *phaC* in CGA2023.

Several of the genes in the RegSR regulon are likely more important under aerobic growth conditions. For instance, two genes down-regulated in CGA2023, *rpa0549* and *rpa0550*, encode homologs of the anti-sigma factor ChrR and the sigma factor, RpoE, respectively, which play a role in oxidative stress caused by singlet oxygen (45–48). Additionally, a third of the remaining genes encode proteins involved in inorganic ion transport (Table S1). As shown in Table S1, many of these genes are down-regulated in CGA2023 and include genes involved in Fe(III) uptake such as *fbpA, fbpB*, *exbB3* and *exbD3*. Uptake of Fe(III) is important under aerobic conditions since most Fe(II) is oxidized to Fe(III) under these conditions, and we found CGA2023 grew more slowly than CGA010 under aerobic, iron-limiting conditions (Fig. S5). Growth was not significantly different under anaerobic, iron-limited conditions (Fig. S5).

## Discussion

Expression of *pioABC* is a poor marker for anoxygenic phototrophic Fe(II) oxidation in the environment since expression of *pioABC* occurs under photoheterotrophic conditions and photoautotrophic conditions with H_2_ and thiosulfate (36, 49). It was proposed that *pioABC* expression could be a marker for photoautotrophic conditions since high levels of expression were only observed under photoautotrophic conditions when compared to photoheterotrophic growth with acetate (49). However, we found that expression of *pioABC* under photoheterotrophic conditions is dependent on the oxidation state of the carbon source provided and whether cells are carrying out nitrogen fixation. Expression of *pioABC* was highest when cells were grown on malate and carrying out nitrogen fixation and lowest when cells were grown with a more reduced carbon substrate such as acetate or benzoate and not fixing nitrogen. Malate is the most oxidized carbon source used in this study and nitrogen fixation requires large amounts of reducing equivalents. This suggests that expression of *pioABC* is regulated in response to the cellular redox state, and *pioABC* expression may be indicative of conditions in which there is an increased cellular demand for reducing equivalents.

We also found that increased expression of *pioABC* under these conditions requires the redox-sensing, two-component system RegSR. Under aerobic conditions, the RegS homolog, RegB, senses intracellular redox status using a conserved cytoplasmic cysteine residue. This cysteine can either form an intramolecular disulfide bond or can be modified by sulfenic acid, repressing kinase activity of RegB (50, 51). RegB also interacts with quinones in the cytoplasmic membrane through a conserved quinone-binding site to monitor the redox state of the quinone pool and alter gene expression in response (39, 52). Since *pioABC* is only expressed under microaerobic or anaerobic conditions, we believe that RegSR is regulating *pioABC* expression in response to changes in the redox state of the quinone pool. It is still unclear if the transcription regulator RegR can bind to the promoter of *pioABC*, and what role phosphorylation of RegR plays in regulating expression of *pioABC*. Interestingly, RegBA homologs have been implicated in regulating Fe(II) oxidation in the acidophilic Fe(II)-oxidizer, *Acidithiobacillus ferrooxidans*, which suggests that this redox-sensing pathway may have been co-opted to regulate other Fe(II)-oxidizing systems (53).

Since electrons from Fe(II) oxidation feed into the quinone pool and are used for reverse electron transfer to produce NADH, we present a model in which RegSR controls *pioABC* expression in response to the redox state of the quinone pool (Fig. 6). RegSR could provide feedback regulation that would allow the cell to up-regulate expression of *pioABC* when there is an increased demand for reducing equivalents such as during photoautotrophic growth and photoheterotrophic growth on an oxidized carbon substrate. It could also down-regulate expression of *pioABC* to prevent the cells from becoming overly reduced under photoheterotrophic conditions on more reduced carbon substrates (Fig. 6). Regulation of *pioABC* expression by the redox state of the quinone pool may explain why a lower rate of Fe(II) oxidation was observed for *R. palustris* TIE-1 when both Fe(II) and H_2_ are present under photoautotrophic conditions versus Fe(II) alone, and why Fe(II) oxidation under photoheterotrophic conditions by *R. palustris* TIE-1 is dependent on the carbon substrate provided (13, 54).

**Figure 6.**
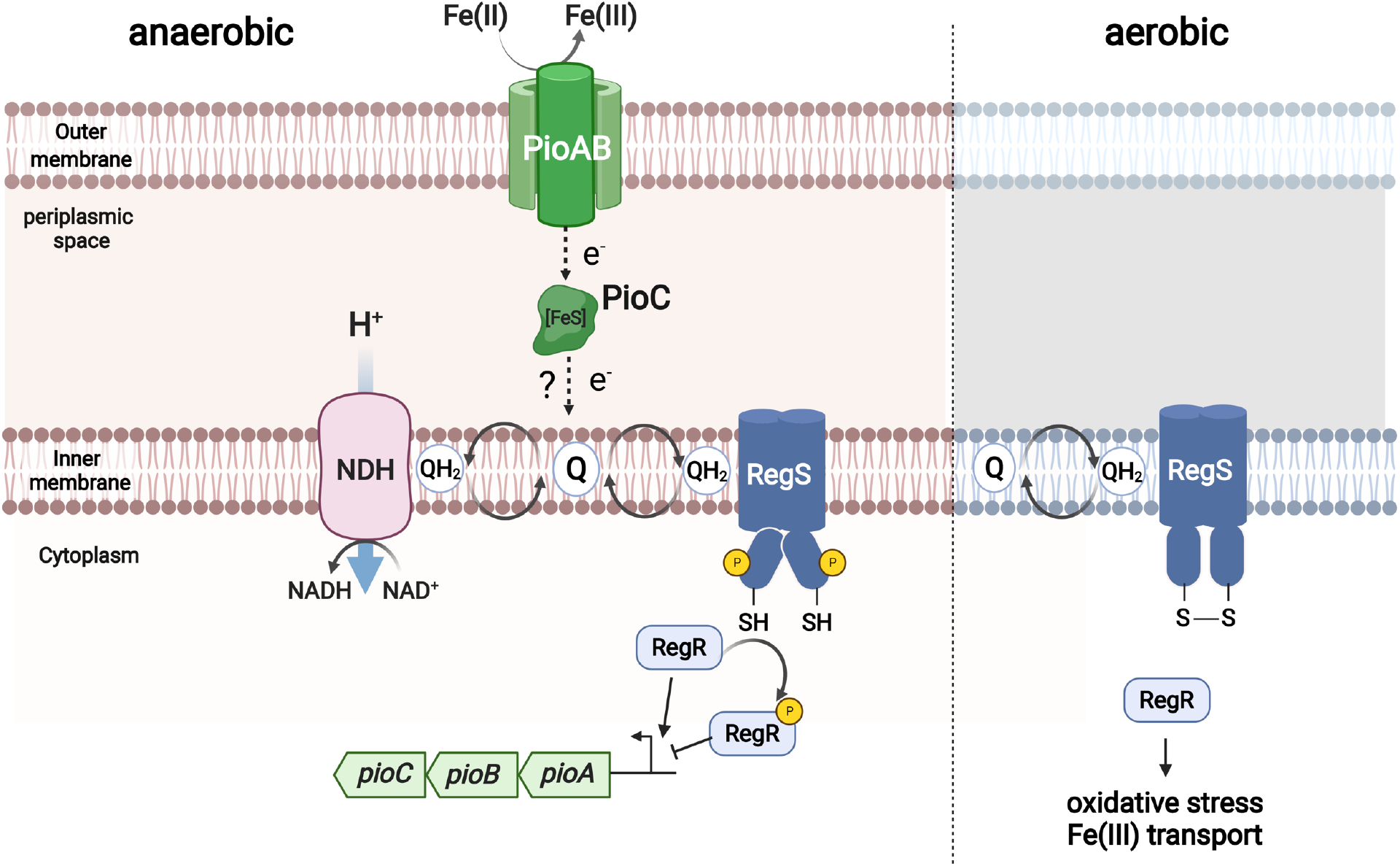
Regulation of *pioABC* by RegSR. PioABC forms an electron transfer pathway that carries electrons from oxidation of Fe(II) to the quinone pool (dotted line), where the electrons enter the quinone pool and are used by NADH dehydrogenase (NDH) to reduce NAD^+^. It is still unclear how electrons enter the quinone pool (?). RegS is active when it binds ubiquinol, and a cysteine in its cytoplasmic domain is reduced. Under aerobic conditions, RegS is largely unphosphorylated because the cysteine is oxidized and the quinone pool is more oxidized. Under anaerobic conditions, the redox-sensitive cysteine residue in RegS is reduced, and the phosphorylation state of RegS is determined by the redox state of the quinone pool. This would allow RegSR to modulate expression of *pioABC* based on the redox state of the quinone pool, leading to increased expression when the quinone pool is more oxidized and decreased expression when the quinone pool is more reduced. Q, quinone; QH_2_, reduced quinone. Image created with BioRender.com.

*R. palustris* CGA010 and the PNSB *Rhodomicrobium vannielii* encode *pioABC* but are unable to grow photoautotrophically using Fe(II) as the electron donor (16, 32). It was unclear why these organisms encode *pioABC,* but one possibility was that Fe(II) oxidation plays a role in ameliorating Fe(II) toxicity (25). However, we saw no difference in growth when cells were grown with a high concentration of Fe(II) and unable to carry out Fe(II) oxidation versus cells grown at a lower concentration of Fe(II) (Fig. S3). This indicates that Fe(II) oxidation is not required for dealing with Fe(II) toxicity in *R. palustris* CGA010. We think the more likely explanation, at least for *R. palustris* CGA010, is that Fe(II) oxidation is playing a role as an electron donor in photolithoheterotrophic growth. Even on oxidized carbon substrates, a significant portion of electron carriers must be oxidized by the Calvin cycle in *R. palustris* to recycle electron carriers and maintain redox balance (55, 56). This means that some of the carbon substrate is not incorporated into biomass but is instead oxidized to produce reducing equivalents and CO_2_ through the TCA cycle (55, 56). Fe(II) oxidation may give the cell access to an additional source of electrons, allowing the cell to incorporate more of the carbon substrate into biomass. However, the cell would still be limited by its ability to oxidize the reducing equivalents it generates. This may explain why a higher cell yield was observed under nitrogen-fixing conditions. Additional CO_2_ was not provided to the cultures, which means the ability of the cells to oxidize reducing equivalents using CO_2_ fixation was likely limited. Nitrogen fixation can also be used to recycle electron carriers, and H_2_ production by nitrogenase can replace CO_2_ fixation to maintain redox balance in *R. palustris* (55, 56). By growing the cells under nitrogen-fixing conditions, the cells can recycle electron carriers without the need to fix CO_2_.

Fe(II) oxidation is not restricted to photoautotrophic growth in anoxygenic phototrophs, and we have shown that Fe(II) oxidation and the proteins required for Fe(II) oxidation can play a role in photolithoheterotrophic metabolism in at least one anoxygenic phototroph. The ability to obtain electrons from both inorganic and organic electron donors could provide an advantage to anoxygenic phototrophs in heterogenous environments. More work is needed to understand how prevalent photolithoheterotrophy is with Fe(II)-oxidizing anoxygenic phototrophs and where this type of metabolism is likely to occur. This study indicates that Fe(II) oxidation in aquatic environments is influenced not only by the amount of Fe(II) present but is also likely influenced by organic substrate and nitrogen availability.

## Materials and methods

### Reagents, strains, and culture methods

All strains and plasmids used are listed in Table S2. *R. palustris* strains were grown anaerobically in sealed Hungate tubes with a headspace of nitrogen gas and carbon sources were added after autoclaving from sterile, anaerobic stock solutions. Each tube contained 10 mL of either defined mineral minimal medium (PM) or nitrogen-fixing medium (NFM), which is PM medium without ammonium sulfate added (Kim and Hardwood, 1991). All liquid growth medium was deaerated by bubbling argon gas through the medium and dispensing the liquid into Hungate tubes in an anaerobic glove box. Media was adjusted to appropriate pH using NaOH or HCl when noted. All *R. palustris* cultures were grown at 30°C under high light (30 μmol photons/m^2^/s) provided by a 60 W incandescent light bulb (General Electric, Boston, MA, USA). Strains were initially grown in PM medium supplemented with 20 mM acetate and 0.1% yeast extract, then diluted into either PM or NFM medium including the stated carbon sources which were added to a final concentration of 40 mM carbon or 20 mM where stated. These cultures were used to inoculate cultures in triplicate for subsequent growth analysis. To grow *R. palustris* using hydrogen gas as an electron donor, PM medium was prepared as before, however the headspace of each tube was flushed with 100% H_2_ for 45 seconds after autoclaving. For photoautotrophic conditions, sodium bicarbonate was added to final concentrations of 10 mM. Fe(II) oxidation conditions were created by the addition of 2.5 mM ferrous chloride to the medium, which formed a precipitate that could be removed with the addition of 2.5 mM nitriloacetic acid. Protein yields were similar with or without the nitriloacetic acid. Optical density readings were taken using a Genesys 50 UV-visible spectrophotometer, and total protein concentrations were determined using the Bio-Rad Protein assay kit. *R. palustris* was also grown aerobically in 250 mL open-mouthed flasks containing 50 mL of PM medium that was not deaerated with argon. Cultures were grown with no direct illumination at 30°C and shaking at 250 rpm. Optical density readings were recorded by sampling one milliliter of culture at each time point. *Escherichia coli* strains were grown in Lysogeny Broth (LB) medium at 37°C. When necessary, *R. palustris* and *E. coli* cultures were supplemented with gentamicin (Gm) at a concentration of 100 μg/mL and 20 μg/mL, respectively.

### Genetic manipulation of *R. palustris*

All strains and plasmids are listed in Table S2. Details of strain and plasmid construction can be found in SI Materials and Methods.

### RNA extraction, sequencing, and gene expression analysis

*R. palustris* strains were grown to an optical density of 0.45-0.47 at 660 nm, at which point 10 mL of cells were harvested from each replicate by centrifugation at 4,000 x g for 10 minutes at 4°C. The supernatant was removed, and cells were immediately frozen using liquid nitrogen followed by storage at −80°C until further analysis. Additional details on RNA extraction, sequencing, and gene expression analysis can be found in the SI Materials and Methods.

### Data accession number

All transcriptomic data are available as raw sequencing reads deposited in the NCBI Gene Expression Omnibus under the accession number GSE150608.

### Iron-oxidation assay

Fe(II) oxidation assays were performed in 25 mL Balch tubes containing 5 mL of Fe(II) oxidation medium consisting of (per liter) 0.46 g of NaCl, 0.117 g of MgSO4.7H_2_O, 0.42 g of NaHCO_3_, 0.8 g of Na_3_C_6_H_5_O_7_ and 20 g of HEPES buffer. pH of the medium was adjusted to 7. The Balch tubes containing the medium were made anaerobic by sparging with N_2_: CO_2_ (80:20) gas mixture followed by sealing with butyl rubber stoppers and aluminum crimps. Fe(II) oxidation assays were performed using the cells grown photo-autotrophically on thiosulfate as the electron donor. Cells were pelleted by centrifuging at 16, 000 × g for 3 minutes followed by washing with the Fe(II) oxidation medium and resuspension in the same medium. Resuspended cells were added to the Balch tubes anaerobically to achieve final cell concentrations of 2×10^9^cells/mL. Filtered ferrous chloride solution was added anaerobically to achieve final Fe(II) concentrations of approximately 1 mM. Tubes were incubated at room temperature in front of a 60 W full spectrum bulb. 100 μL of the samples were periodically collected anaerobically and added to 900 μL of 0.5 N HCl. Fe(II) in the samples was quantified using the ferrozine assay (57).

## Supporting information

All supplemental materials

## Acknowledgements

Support for this work was provided by the US. Department of Energy, Office of Science, Basic Energy Sciences, Physical Biosciences program (award number DE-FG-02-05ER15707).

## References

1. A. Eiler, Evidence for the ubiquity of mixotrophic bacteria in the upper ocean: implications and consequences. Appl. Environ. Microbiol. 72, 7431–7437 (2006).

2. M. Hartmann, et al., Mixotrophic basis of Atlantic oligotrophic ecosystems. Proc. Natl. Acad. Sci. U.S.A. 109, 5756–5760 (2012).

3. M. C. Muñoz-Marín, et al., Mixotrophy in marine picocyanobacteria: use of organic compounds by *Prochlorococcus* and *Synechococcus*. ISME J. 14, 1065–1073 (2020).

4. J. Tittel, et al., Mixotrophs combine resource use to outcompete specialists: implications for aquatic food webs. Proc. Natl. Acad. Sci. U.S.A. 100, 12776–12781 (2003).

5. A.-M. Bruun, K. Finster, H. P. Gunnlaugsson, P. N⊘rnberg, M. W. Friedrich, A comprehensive investigation on iron cycling in a freshwater seep including microscopy, cultivation and molecular community analysis. Geomicrobiol. J. 27, 15–34 (2010).

6. D. Emerson, Biogeochemistry and microbiology of microaerobic Fe(II) oxidation. Biochem. Soc. Trans. 40, 1211–1216 (2012).

7. D. Emerson, J. V. Weiss, Bacterial iron oxidation in circumneutral freshwater habitats: findings from the field and the laboratory. Geomicrobiol. J. 21, 405–414 (2004).

8. D. Emerson, E. J. Fleming, J. M. McBeth, Iron-Oxidizing bacteria: an environmental and genomic perspective. Annu. Rev. Microbiol. 64, 561–583 (2010).

9. C. Bryce, et al., Microbial anaerobic Fe(II) oxidation – ecology, mechanisms and environmental implications. Environ. Microbiol. 20, 3462–3483 (2018).

10. A. Kappler, et al., An evolving view on biogeochemical cycling of iron. Nat Rev Microbiol 19, 360–374 (2021).

11. A. Jain, J. A. Gralnick, Engineering lithoheterotrophy in an obligate chemolithoautotrophic Fe(II) oxidizing bacterium. Sci Rep 11, 2165 (2021).

12. E. M. Muehe, S. Gerhardt, B. Schink, A. Kappler, Ecophysiology and the energetic benefit of mixotrophic Fe(II) oxidation by various strains of nitrate-reducing bacteria. FEMS Microbiol. Ecol. 70, 335–343 (2009).

13. E. D. Melton, C. Schmidt, S. Behrens, B. Schink, A. Kappler, Metabolic flexibility and substrate preference by the Fe(II)-oxidizing purple non-sulphur bacterium *Rhodopseudomonas palustris* strain TIE-1. Geomicrobiol. J. 31, 835–843 (2014).

14. S. H. Kopf, D. K. Newman, Photomixotrophic growth of *Rhodobacter capsulatus* SB1003 on ferrous iron: photomixotrophic iron oxidation. Geobiology 10, 216–222 (2012).

15. J. Jamieson, et al., Identifying and quantifying the intermediate processes during nitrate-dependent iron(II) oxidation. Environ. Sci. Technol. 52, 5771–5781 (2018).

16. S. Heising, B. Schink, Phototrophic oxidation of ferrous iron by a *Rhodomicrobium vannielii* strain. Microbiology 144, 2263–2269 (1998).

17. A. Chakraborty, E. E. Roden, J. Schieber, F. Picardal, Enhanced growth of *Acidovorax* sp. strain 2AN during nitrate-dependent Fe(II) oxidation in batch and continuous-flow systems. Appl. Environ. Microbiol. 77, 8548–8556 (2011).

18. N. C. Caiazza, D. P. Lies, D. K. Newman, Phototrophic Fe(II) oxidation promotes organic carbon acquisition by *Rhodobacter capsulatus* SB1003. Appl. Environ. Microbiol. 73, 6150–6158 (2007).

19. F. Widdel, et al., Ferrous iron oxidation by anoxygenic phototrophic bacteria. Nature 362, 834–836 (1993).

20. A. Ehrenreich, F. Widdel, Anaerobic oxidation of ferrous iron by purple bacteria, a new type of phototrophic metabolism. Appl. Environ. Microbiol. 60, 4517–4526 (1994).

21. Y. Jiao, A. Kappler, L. R. Croal, D. K. Newman, Isolation and characterization of a genetically tractable photoautotrophic Fe(II)-oxidizing bacterium, *Rhodopseudomonas palustris* strain TIE-1. Appl. Environ. Microbiol. 71, 4487–4496 (2005).

22. S. Heising, L. Richter, W. Ludwig, B. Schink, *Chlorobium ferrooxidans* sp. nov., a phototrophic green sulfur bacterium that oxidizes ferrous iron in coculture with a *Geospirillum* sp. strain. Arch. Microbiol. 172, 116–124 (1999).

23. K. Laufer, et al., Physiological characterization of a halotolerant anoxygenic phototrophic Fe(II)-oxidizing green-sulfur bacterium isolated from a marine sediment. FEMS Microbiol. Ecol. 93, fix054 (2017).

24. K. L. Straub, F. A. Rainey, F. Widdel, *Rhodovulum iodosum* sp. nov. and *Rhodovulum robiginosum* sp. nov., two new marine phototrophic ferrous-iron-oxidizing purple bacteria. Int. J. Syst. Evol. Microbiol. 49, 729–735 (1999).

25. A. J. Poulain, D. K. Newman, *Rhodobacter capsulatus* catalyzes light-dependent Fe(II) oxidation under anaerobic conditions as a potential detoxification mechanism. Appl. Environ. Microbiol. 75, 6639–6646 (2009).

26. L. R. Croal, C. M. Johnson, B. L. Beard, D. K. Newman, Iron isotope fractionation by Fe(II)-oxidizing photoautotrophic bacteria. Geochim. Cosmochim. Acta 68, 1227–1242 (2004).

27. Y. Oda, et al., Multiple genome sequences reveal adaptations of a phototrophic bacterium to sediment microenvironments. Proc. Natl. Acad. Sci. U.S.A. 105, 18543–18548 (2008).

28. F. W. Larimer, et al., Complete genome sequence of the metabolically versatile photosynthetic bacterium *Rhodopseudomonas palustris*. Nat. Biotechnol. 22, 55–61 (2004).

29. F. E. Rey, Y. Oda, C. S. Harwood, Regulation of uptake hydrogenase and effects of hydrogen utilization on gene expression in *Rhodopseudomonas palustris*. J. Bacteriol. 188, 6143–6152 (2006).

30. J. J. Huang, E. K. Heiniger, J. B. McKinlay, C. S. Harwood, Production of hydrogen gas from light and the inorganic electron donor thiosulfate by *Rhodopseudomonas palustris*. Appl. Environ. Microbiol. 76, 7717–7722 (2010).

31. J. P. Rolls, E. S. Lindstrom, Effect of thiosulfate on the photosynthetic growth of *Rhodopseudomonas palustris*. J. Bacteriol. 94, 860–866 (1967).

32. Y. Jiao, D. K. Newman, The *pio* operon is essential for phototrophic Fe(II) oxidation in *Rhodopseudomonas palustris* TIE-1. J. Bacteriol. 189, 1765–1773 (2007).

33. D. Gupta, et al., Photoferrotrophs produce a PioAB electron conduit for extracellular electron uptake. mBio 10, e02668–19 (2019).

34. L. J. Bird, et al., Nonredundant roles for cytochrome *c*2 and two high-potential iron-sulfur proteins in the photoferrotroph *Rhodopseudomonas palustris* TIE-1. J. Bacteriol. 196, 850–858 (2014).

35. L. J. Bird, V. Bonnefoy, D. K. Newman, Bioenergetic challenges of microbial iron metabolisms. Trends Microbiol. 19, 330–340 (2011).

36. A. Bose, D. K. Newman, Regulation of the phototrophic iron oxidation (*pio*) genes in *Rhodopseudomonas palustris* TIE-1 is mediated by the global regulator, FixK. Mol. Microbiol. 79, 63–75 (2011).

37. F. E. Rey, C. S. Harwood, FixK, a global regulator of microaerobic growth, controls photosynthesis in *Rhodopseudomonas palustris*. Mol. Microbiol. 75, 1007–1020 (2010).

38. M. W. Sganga, C. E. Bauer, Regulatory factors controlling photosynthetic reaction center and light-harvesting gene expression in *Rhodobacter capsulatus*. Cell 68, 945–954 (1992).

39. L. R. Swem, X. Gong, C.-A. Yu, C. E. Bauer, Identification of a ubiquinone-binding site that affects autophosphorylation of the sensor kinase RegB. J. Biol. Chem. 281, 6768–6775 (2006).

40. J. E. Kumka, H. Schindel, M. Fang, S. Zappa, C. E. Bauer, Transcriptomic analysis of aerobic respiratory and anaerobic photosynthetic states in Rhodobacter capsulatus and their modulation by global redox regulators RegA, FnrL and CrtJ. Microb. Genom. 3, e000125 (2017).

41. S. Elsen, L. R. Swem, D. L. Swem, C. E. Bauer, RegB/RegA, a highly conserved redox-responding global two-component regulatory system. Microbiol. Mol. Biol. Rev. 68, 263–279 (2004).

42. J. Wu, C. E. Bauer, “RegB/RegA, a global redox-responding two-component system” in Bacterial Signal Transduction: Networks and Drug Targets, Advances in Experimental Medicine and Biology., R. Utsumi, Ed. (Springer New York, 2008), pp. 131–148.

43. B. C. Kelley, C. M. Meyer, C. Gandy, P. M. Vignais, Hydrogen recycling by *Rhodopseudomonas capsulata*. FEBS Lett. 81, 281–285 (1977).

44. R. T. Haug, “Metabolic and nutritional classifications” in Lessons in Environmental Microbiology, 1st Ed. (CRC Press, 2019), pp. 139–169.

45. J. R. Anthony, K. L. Warczak, T. J. Donohue, A transcriptional response to singlet oxygen, a toxic byproduct of photosynthesis. Proc. Natl. Acad. Sci. U.S.A. 102, 6502–6507 (2005).

46. T.-W. Nam, E. C. Ziegelhoffer, R. A. S. Lemke, T. J. Donohue, Proteins needed to activate a transcriptional response to the reactive oxygen species singlet oxygen. mBio 4, e00541–12 (2013).

47. J. Glaeser, M. Zobawa, F. Lottspeich, G. Klug, Protein synthesis patterns reveal a complex regulatory response to singlet oxygen in *Rhodobacter*. J. Proteome Res. 6, 2460–2471 (2007).

48. A. M. Nuss, et al., DegS and RseP homologous proteases are involved in singlet oxygen dependent activation of RpoE in *Rhodobacter sphaeroides*. PLoS ONE 8, e79520 (2013).

49. C. Bryce, et al., Proteome response of a metabolically flexible anoxygenic phototroph to Fe(II) Oxidation. Appl. Environ. Microbiol. 84, e01166–18 (2018).

50. L. R. Swem, Signal transduction by the global regulator RegB is mediated by a redox-active cysteine. EMBO J. 22, 4699–4708 (2003).

51. J. Wu, et al., RegB kinase activity is repressed by oxidative formation of cysteine sulfenic acid. J. Biol. Chem. 288, 4755–4762 (2013).

52. J. Wu, C. E. Bauer, RegB kinase activity is controlled in part by monitoring the ratio of oxidized to reduced ubiquinones in the ubiquinone pool. mBio 1, e00272–10 (2010).

53. D. Moinier, D. Byrne, A. Amouric, V. Bonnefoy, The global redox responding RegB/RegA signal transduction system regulates the genes involved in ferrous iron and inorganic sulfur compound oxidation of the acidophilic *Acidithiobacillus ferrooxidans*. Front. Microbiol. 8, 1277 (2017).

54. L. R. Croal, Y. Jiao, A. Kappler, D. K. Newman, Phototrophic Fe(II) oxidation in an atmosphere of H_2_ : implications for Archean banded iron formations. Geobiology 7, 21–24 (2009).

55. J. B. McKinlay, C. S. Harwood, Carbon dioxide fixation as a central redox cofactor recycling mechanism in bacteria. Proc. Natl. Acad. Sci. U.S.A. 107, 11669–11675 (2010).

56. J. B. McKinlay, C. S. Harwood, Calvin cycle flux, pathway constraints, and substrate oxidation state together determine the H_2_ biofuel yield in photoheterotrophic bacteria. mBio 2, e00323–10 (2011).

57. L. L. Stookey, Ferrozine---a new spectrophotometric reagent for iron. Anal. Chem. 42, 779–781 (1970).

