## Supplementary material for "Iron oxidation is regulated by the two-component system, RegSR, and plays a role in photolithoheterotrophic growth in *Rhodopseudomonas palustris*": All supplemental materials

**This PDF file includes:**

Supplemental text
Figures S1 to S5
Table S1 to S2
SI references

**Supplementary Information Text**

**SI Material and Methods**

*Genetic manipulation of* R. palustris. The *regSR* operon was expressed *in trans* from plasmid pBBPgdh (Table S2). Genomic DNA from *R. palustris* CGA010 was used as template during PCR with Phusion High-Fidelity DNA polymerase (NEB, Ipswich, MA, USA). A ~2.4-kb PCR fragment was amplified containing the *regSR* operon plus 200 bp upstream of the *regS* start codon and 200 bp downstream of the *regR* stop codon. All PCR fragments were purified using the QIAquick PCR Purification Kit (QIAGEN, Hilden, Germany). These fragments were incorporated into the multiple cloning site of plasmid pBBPgdh directly downstream of the GAPDH promoter via *E. coli* DH5α-mediated assembly as described previously (1). Correct location and orientation of the fragment with respect to the promoter was verified with PCR and sequencing. The assembled p-*regSR* plasmid and pBBPgdh were mobilized into *R. palustris* CGA2023 by conjugation with *E. coli* S17-1 and successful conjugates were selected for on PM agar supplemented with gentamicin and 10 mM succinate.

In-frame deletions of *pioA* and *pioABC* were created by PCR using Phusion High-Fidelity DNA polymerase (New England Biolabs) to amplify a 1-kb fragment upstream of the coding region and 1-kb downstream of the stop codon from purified *R. palustris* CGA010 genomic DNA. The suicide vector pJQ200SK was amplified using PrimeSTAR® Max DNA polymerase (Takara Bio). All fragments were assembled as described in (1). The assembled plasmids, pJQ-Δ*pioA* and pJQ-Δ*pioABC*, were mobilized into *R. palustris* by conjugation with *E. coli* S17-1, and double-crossover events for deletions or allelic exchange was achieved using a selection and screening strategy described previously (2). All deletions were verified by PCR and sequencing of the resulting PCR product.

*RNA Extraction and Sequencing.* Cell pellets were resuspended in 1 mL QIAzol Lysis Reagent (QIAGEN, Hilden, Germany) and transferred into a 2 mL screw-cap tube containing 0.1 mm diameter zirconia/silicate beads (BioSpec Products INC., Bartlesville, OK, USA). Each sample was homogenized in a Mini-BeadBeater-24 (BioSpec Products INC., Bartlesville, OK, USA) at maximum rpm for one minute at 4˚C. Tubes were allowed to cool on ice for at least one minute and the process was repeated four additional times. Cellular debris was removed from the homogenized cell lysate via chloroform extraction and total RNA was isolated using the miRNAeasy Mini Kit (QIAGEN, Hilden, Germany). To remove DNA, each sample was incubated with TURBO DNase (Invitrogen, Carlsbad, CA, USA). The RNeasy MinElute Cleanup Kit (QIAGEN, Hilden, Germany) was used to purify and concentrate total RNA.

Ribosomal RNA depletion, cDNA library construction and sequencing reactions were performed at GENEWIZ, LLC (South Plainfield, NJ, USA). Briefly, RNA samples were quantified and checked for integrity, followed by rRNA depletion using the Ribo-Zero rRNA Removal Kit (Illumina, San Diego, CA, USA). Preparation of cDNA sequencing libraries utilized the NEBNext Ultra RNA Library Prep Kit (NEB, Ipswich, MA, USA). All sequencing reactions, image analysis and base calling were performed on an Illumina HiSeq 2500 instrument (Illumina, San Diego, CA, USA).

*Differential Gene Expression Analysis.* Sequencing data were analyzed using the FastQC application v0.11.8 (https://www.bioinformatics.babraham.ac.uk/projects/fastqc/) to verify high-quality base calling. The TrimGalore script v0.6.2 (https://www.bioinformatics.babraham.ac.uk/projects/trim_galore/) was used to trim adapter sequences, process, and validate reads according to default parameters. All subsequent data processing and analysis was performed on the Avadis software package v3.1.1 (Strand Life Sciences, Bengaluru, India). Trimmed reads were aligned to the published genome sequence of CGA009 (NC_005296). The DESeq2 package in R v3.6 (3) was used with default parameters to determine differentially expressed genes. Genes that exhibited a fold-change ratio greater than or equal to two between CGA010 versus CGA2023 and had a *P*-value less than or equal to 0.05 as determined by DESeq were considered differentially expressed. These data have been deposited in the NCBI Gene Expression Omnibus under the GEO accession number GSE150608.

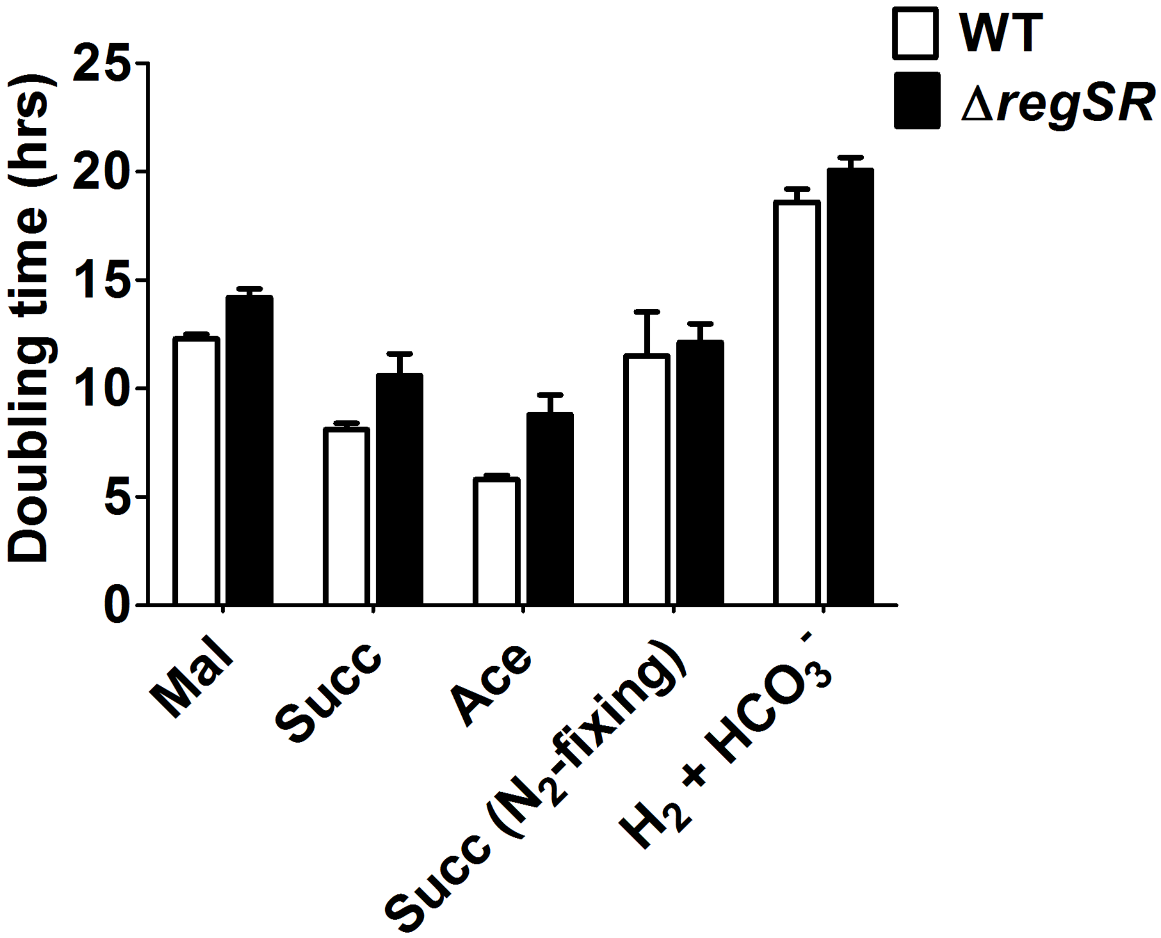


**Figure S1. Doubling times for CGA010 (WT) and CGA2023 (Δ*regSR*) grown under different conditions**. Cells were grown in minimal medium with ammonium sulfate or without ammonium sulfate ($N_2$-fixing). All carbon substrates were added to a final concentration of 40 mM. Data represent average of three replicates, and error bars represent standard deviation. Mal, photoheterotrophic growth with malate; Succ, photoheterotrophic growth with succinate; Ace, photoheterotrophic growth with acetate; $H_2$ + $HCO_3^-$, photoautotrophic growth with hydrogen and bicarbonate.

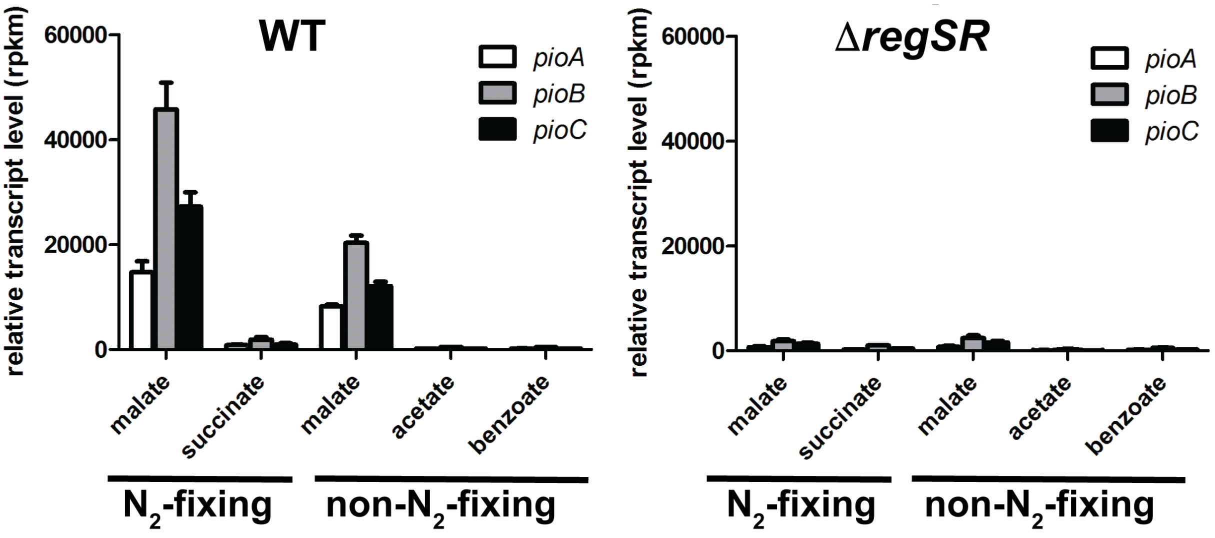


**Figure S2. Expression of *pioABC* requires RegSR and is affected by both the oxidation state of the carbon substrate and nitrogen fixation under photoheterotrophic conditions.** Shown is the relative *pioABC* transcript levels expressed as reads per kilobase per million reads (rpkm) in *Rps. palustris* CGA010 (WT) and CGA2023 (Δ*regSR*) as determined by RNA-seq analysis. Cells were grown photoheterotrophically using the indicated carbon substrate, supplied at a final concentration of 40 mM carbon, with (non-$N_2$-fixing) or without ($N_2$-fixing) ammonium sulphate. Carbon substrates are ordered by decreasing oxidation state. Data are averages from duplicate cultures and the bars show SEM.

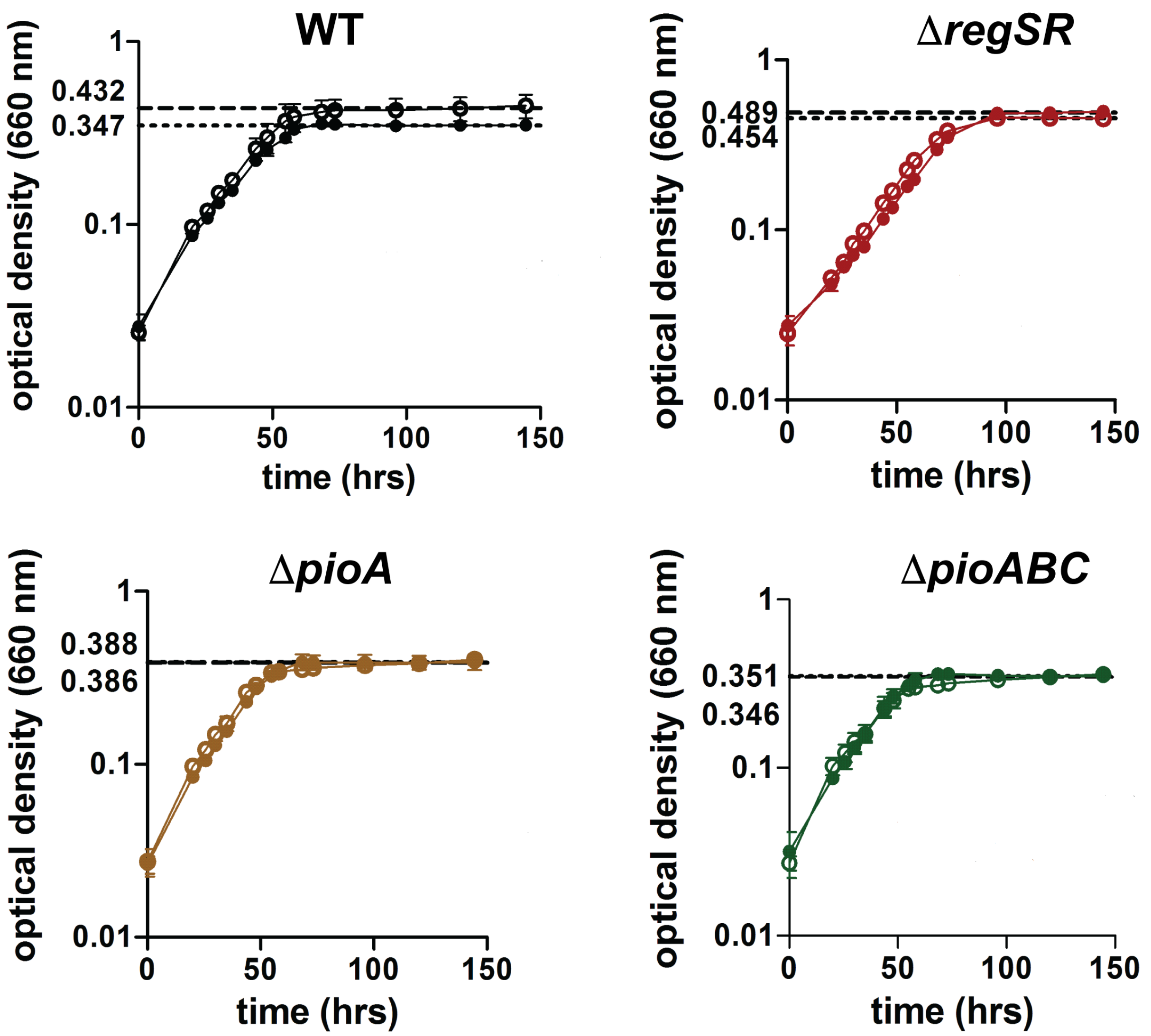


**Figure S3. *R. palustris* cell density increases under Fe(II)-oxidizing conditions.** Growth of *R. palustris* CGA010 (WT) and CGA2023 (Δ*regSR),* Δ*pioA*, and Δ*pioABC* strains in minimal medium without ($N_2$-fixing) ammonium sulfate, 5 mM malate, and with (open circles) or without 2.5 mM $FeCl_2$ (filled circles) was determined using optical density at 660 nm. Bold dotted line indicates final optical density reach with 2.5 mM $FeCl_2$ added, and dotted line represents final optical density reached without $FeCl_2$ added. Data represent the average of three replicates and the error bars represent standard deviation.

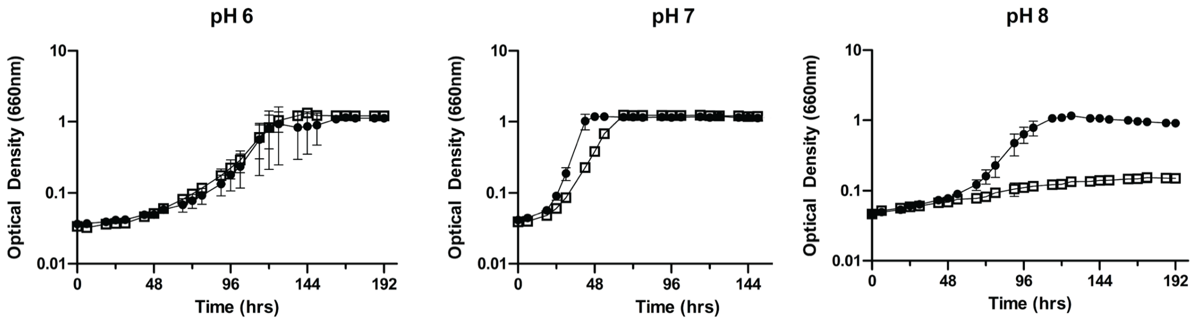


**Figure S4.** ***R. palustris*** **cells lacking an intact** ***regSR*** **operon are more sensitive to high pH.** Photoheterotrophic growth of *R. palustris* CGA010 (black circles) and CGA2023 (open squares) on 10mM succinate at varying pH values. Growth rates were similar at pH values of 6 and 7, while growth of CGA2023 was inhibited at pH 8. All values are the average of three replicates and the error bars represent standard deviation.

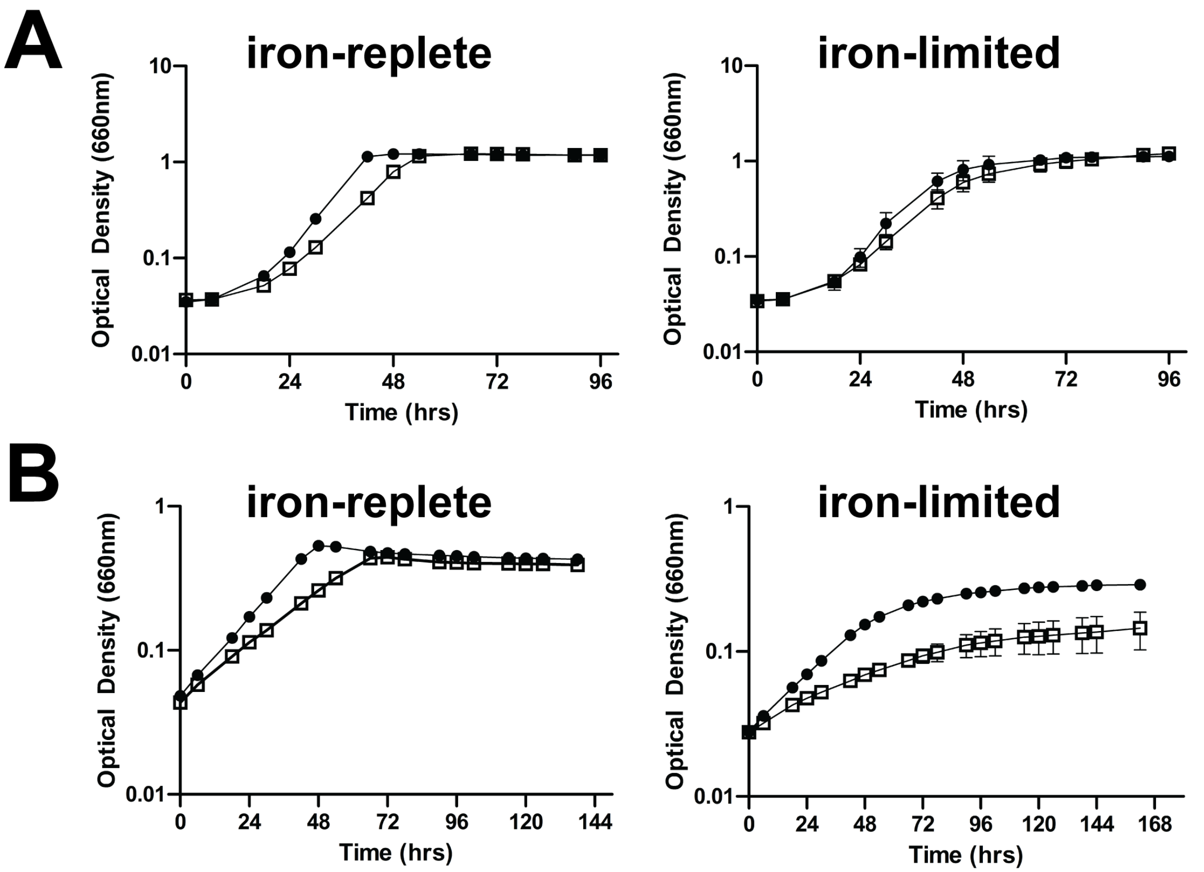


**Figure S5. CGA2023 shows a growth defect in iron-limited conditions in the presence of oxygen but not in the absence of oxygen.** Photoheterotrophic growth (A) and aerobic growth (B) of CGA010 (black circles) and CGA2023 (open squares) in iron replete or iron-limited minimal medium with 10 mM succinate. Growth was determined by measuring optical density at 660 nm. Data represent the average of three replicates and the error bars represent standard deviation.

**Table S1. Genes differentially expressed under all conditions tested.**

| Locus tag [Gene name] | Description | Fold-change ratio (CGA2023 vs. CGA010)[a] | | | | |
|---|---|---|---|---|---|---|
| | | non-$N_2$-fixing | | | $N_2$-fixing | |
| | | Mal | Ace | Benz | Mal | Suc |
| RPA0198 | possible acetate transporter, stationary-phase anti-death (SAD) family | -9.7 | -6.4 | -13.8 | -9.3 | -12.2 |
| RPA0549 [*chrR*] | cytochrome cycA regulator | -5.2 | -2.8 | -3.4 | -3.2 | -3.9 |
| RPA0550 | RNA polymerase ECF-type sigma factor | -20.1 | -15.2 | -16.2 | -9.3 | -28.6 |
| RPA0566 | conserved unknown protein | 2.4 | 5.5 | 33.4 | 2.4 | 2.5 |
| RPA0766 | transcriptional regulator, Crp/Fnr family | -3.8 | -3.2 | -2.2 | -4.7 | -5.4 |
| RPA0787 | putative heat shock protein (htpX) | -5.0 | -5.4 | -8.1 | -4.4 | -6.7 |
| RPA0882 | ABC transporter, permease | 2.1 | 2.5 | 2.0 | 2.7 | 2.8 |
| RPA0883 | ABC transporter, ATP-binding protein | 2.1 | 2.5 | 2.5 | 2.9 | 3.2 |
| RPA0884 | ABC transporter, substrate-binding protein | 2.4 | 3.2 | 2.6 | 2.8 | 4.0 |

| | | | | | | |
|---|---|---|---|---|---|---|
| RPA1009 | possible cytochrome P450 | 3.3 | 2.2 | 2.1 | 2.5 | 2.8 |
| RPA1136 [*ictP*] | putative L-lactate permease | 2.1 | 3.9 | 6.9 | 2.8 | 5.7 |
| RPA1259 | putative cation-transporting P-type ATPase | -5.6 | -2.4 | -3.0 | -4.6 | -4.6 |
| RPA1260 | Universal stress protein (Usp) | -5.8 | -2.5 | -2.6 | -5.9 | -5.8 |
| RPA1301 [*pfpI*] | putative intracellular protease | 2.6 | 3.3 | 17.6 | 2.5 | 2.6 |
| RPA1497 [*mexC1*] | putative RND multidrug efflux protein | -4.6 | -17.4 | -6.2 | -6.1 | -15.5 |
| RPA1498 [*mexD2*] | RND multidrug efflux transporter | -4.0 | -14.2 | -7.6 | -3.5 | -10.1 |
| RPA1500 | unknown protein | 4.0 | 2.2 | 11.2 | 3.1 | 2.3 |
| RPA1515 | conserved unknown protein | -17.7 | -14.0 | -26.5 | -14.1 | -22.6 |
| RPA1587 | hypothetical protein | -3.4 | -3.1 | -3.8 | -3.6 | -4.2 |
| RPA1685 | possible serine protease/outer membrane autotransporter | -3.0 | -7.0 | -3.6 | -3.1 | -4.4 |
| RPA1738 | putative branched-chain amino acid transport system ATPase | 3.0 | 3.3 | 3.2 | 2.9 | 4.2 |

| | | | | | | |
|---|---|---|---|---|---|---|
| RPA1739 | putative branched-chain amino acid transport system permease protein | 3.1 | 3.4 | 3.7 | 3.0 | 4.5 |
| RPA1740 | possible branched-chain amino acid transport system permease protein | 2.7 | 3.0 | 3.7 | 2.9 | 4.3 |
| RPA1741 | possible branched-chain amino acid transport system substrate-binding protein | 3.0 | 4.4 | 5.6 | 2.4 | 4.1 |
| RPA1845 | putative tonB-dependent receptor protein | -10.2 | -15.4 | -8.0 | -13.3 | -15.6 |
| RPA1846 | unknown protein | -12.6 | -13.2 | -5.2 | -10.6 | -16.5 |
| RPA1847 | conserved hypothetical protein | -12.0 | -20.8 | -11.2 | -13.7 | -17.2 |
| RPA1848 | Uncharacterized iron-regulated membrane protein DUF337 | -12.0 | -14.8 | -9.4 | -9.6 | -20.0 |
| RPA1849 | conserved hypothetical protein | -8.9 | -13.2 | -8.3 | -8.1 | -21.2 |
| RPA1874 | hypothetical protein | -6.4 | -4.0 | -5.1 | -2.8 | -5.1 |
| RPA1875 | possible uncharacterized iron-regulated membrane protein | -6.8 | -4.4 | -11.6 | -4.0 | -8.2 |

| | | | | | | |
|---|---|---|---|---|---|---|
| RPA1876 | putative TonB-dependent iron siderophore receptor | -5.1 | -3.1 | -6.3 | -5.0 | -6.2 |
| RPA2116 | hypothetical protein | -7.5 | -2.6 | -4.6 | -6.5 | -6.7 |
| RPA2117 | putative flavodoxin | -8.0 | -2.2 | -5.2 | -8.8 | -5.4 |
| RPA2119 | putative permease protein of ABC transporter | -6.6 | -4.0 | -3.4 | -3.6 | -6.5 |
| RPA2120 | putative hemin binding protein | -6.3 | -2.6 | -3.3 | -4.4 | -5.8 |
| RPA2121 | conserved unknown protein | -6.5 | -2.2 | -3.1 | -6.2 | -5.3 |
| RPA2122 | putative oxygen-independent coproporphyrinogen III oxidase | -26.3 | -5.2 | -3.9 | -7.8 | -9.8 |
| RPA2159 | hypothetical protein | -12.6 | -18.1 | -6.2 | -4.7 | -5.9 |
| RPA2299 | transcriptional regulator, LuxR family | -4.3 | -2.5 | -11.1 | -5.1 | -6.7 |
| RPA2300 | transcriptional regulator, LuxR family | -10.7 | -8.2 | -17.5 | -10.4 | -11.4 |
| RPA2307 | possible TonB-dependent receptor precursor | -3.9 | -7.8 | -3.5 | -4.2 | -8.0 |
| RPA2308 | possible periplasmic iron siderophore | -2.0 | -3.5 | -2.3 | -2.8 | -3.8 |

| | | | | | | |
|---|---|---|---|---|---|---|
| | binding protein of ABC transporter | | | | | |
| RPA2309 | putative iron chelatin ABC transporter, permease subunit | -2.0 | -5.6 | -3.5 | -2.3 | -3.1 |
| RPA2310 | putative iron ABC transporter ATP-binding protein | -3.2 | -6.0 | -2.8 | -2.5 | -4.8 |
| RPA2311 | hypothetical protein | -5.3 | -6.3 | -6.0 | -4.2 | -10.5 |
| RPA2312 | hypothetical protein | -5.5 | -6.4 | -8.7 | -5.3 | -11.1 |
| RPA2313 | unknown protein | -6.2 | -6.1 | -9.2 | -7.0 | -9.0 |
| RPA2704 | possible Na+/? Antiporter | -2.3 | -2.4 | -2.9 | -2.3 | -3.2 |
| RPA2788 | possible methyltransferase | -2.6 | -3.3 | -4.0 | -2.3 | -3.3 |
| RPA2789 [*phaA*] | pH adaptation $K^+$ efflux system components | -2.7 | -2.6 | -2.1 | -2.1 | -2.9 |
| RPA2790 [*phaC*] | pH adaptation $K^+$ efflux system component | -3.7 | -3.1 | -2.3 | -2.7 | -3.2 |
| RPA2963 | hypothetical protein | -4.0 | -4.4 | -4.6 | -3.0 | -4.6 |
| RPA3054 | transcriptional regulator, Crp/Fnr family | -8.8 | -5.7 | -5.8 | -9.0 | -10.4 |
| RPA3196 | hypothetical protein | 2.9 | 4.1 | 12.1 | 2.4 | 2.8 |

| | | | | | | |
|---|---|---|---|---|---|---|
| RPA3309 | conserved unknown protein | 3.7 | 2.9 | 25.7 | 3.0 | 2.7 |
| RPA3458 | possible TrapT family, dctP subunit, C4-dicarboxylate periplasmic binding protein | -11.0 | -5.2 | -4.6 | -6.9 | -7.2 |
| RPA3459 | possible TrapT family, fused dctM-Q subunits, C4-dicarboxylate transport | -8.0 | -2.9 | -5.3 | -8.0 | -8.4 |
| RPA3491 [*hflK*] | protease subunit HflK | -2.1 | -2.1 | -2.8 | -2.1 | -3.2 |
| RPA3568 | conserved unknown protein | 3.9 | 2.1 | 17.7 | 3.9 | 2.6 |
| RPA3652 | conserved hypothetical protein | 2.5 | 2.8 | 5.4 | 2.5 | 2.1 |
| RPA3843 [*kefC*] | putative potassium efflux transporter | -3.4 | -4.0 | -2.8 | -3.4 | -4.2 |
| RPA3881 | conserved unknown protein | -16.8 | -13.4 | -16.3 | -16.8 | -22.6 |
| RPA4152 [*fbpA*] | periplasmic iron binding protein | -4.8 | -2.9 | -2.5 | -4.8 | -4.9 |
| RPA4153 [*fpbB*] | iron transport system permease protein | -2.9 | -4.3 | -2.4 | -2.9 | -5.2 |
| RPA4279 | hypothetical protein | -4.6 | -4.9 | -5.8 | -4.6 | -7.0 |

| | | | | | | |
|---|---|---|---|---|---|---|
| RPA4280 | transcription regulator, AraC family | -6.2 | -4.5 | -4.1 | -6.2 | -8.9 |
| RPA4683 | hypothetical protein | -37.7 | -2.7 | -7.0 | -37.7 | -14.2 |

[a]Numbers represent fold-change in transcript levels. Negative values indicate higher expression in CGA010 and positive values indicate higher expression in CGA2023.

**Table S2. Strains and plasmids used.**

| Strain or plasmid | Genotype, phenotype | Reference, origin, or description |
|---|---|---|
| ***R. palustris* strains** | | |
| CGA010 | *hupV* repaired derivative of CGA009; 4 bp inserted in *hupV* | (2) |
| CGA2023 | Δ*regSR* mutant of CGA010 | (2) |
| Δ*pioA* | CGA010 with an in-frame deletion of *pioA* (*rpa0746)* | this study |
| Δ*pioABC* | CGA009 with an in-frame deletion of *pioABC* (*rpa0744-rpa0746)* | this study |
| ***E. coli* strains** | | |
| DH5α | *fhuA2* (*argF-lacZ*)*U169 phoA glnV44 80*(*lacZ*)M15 *gyrA96 recA1 relA1 endA1 thi-1 hsdR17* | New England Biolabs |
| S17-1 | *thi pro hdsR* $hdsM^+$ *recA*; chromosomal insertion of RP4-2 (Tc::Mu Km::Tn7) | (4) |
| **Plasmids** | | |
| pJQ200SK | $Gm^R$, *sacB*; mobilizable suicide vector | (5) |
| pBBPgdh | $Gm^R$; pBBR1MCS-5 with *rpa0944* promoter between KpnI and XhoI sites | (6) |
| pJQ-Δ*pioA* | $Gm^R$; in-frame Δ*pioA* (*rpa0746*) cloned into PstI site of pJQ200SK | this study |
| pJQ-Δ*pioABC* | $Gm^R$; in-frame Δ*pioABC* (*rpa0744-rpa0746*) cloned into PstI site of pJQ200SK | this study |
| p-*regSR* | $Gm^R$; *regSR* cloned into pBBPgdh | this study |

**SI References**


1. M. Kostylev, A. E. Otwell, R. E. Richardson, Y. Suzuki, Cloning should be simple: *Escherichia coli* DH5α-mediated assembly of multiple DNA fragments with short end homologies. *PLoS ONE* **10**, e0137466 (2015).

2. F. E. Rey, Y. Oda, C. S. Harwood, Regulation of uptake hydrogenase and effects of hydrogen utilization on gene expression in *Rhodopseudomonas palustris*. *J. Bacteriol.* **188**, 6143–6152 (2006).

3. M. I. Love, W. Huber, S. Anders, Moderated estimation of fold change and dispersion for RNA-seq data with DESeq2. *Genome Biol.* **15**, 550 (2014).

4. R. Simon, U. Priefer, A. Pühler, A broad host range mobilization system for *in vivo* genetic engineering: transposon mutagenesis in gram-negative bacteria. *Nat. Biotechnol.* **1**, 784–791 (1983).

5. J. Quandt, M. F. Hynes, Versatile suicide vectors which allow direct selection for gene replacement in gram-negative bacteria. *Gene* **127**, 15–21 (1993).

6. J. B. McKinlay, C. S. Harwood, Carbon dioxide fixation as a central redox cofactor recycling mechanism in bacteria. *Proc. Natl. Acad. Sci. U.S.A.* **107**, 11669–11675 (2010).
